# Reduced horseshoe crab abundance and feeding activity beneath intertidal oyster aquaculture structures in the Delaware Bay

**DOI:** 10.1101/2020.06.20.162693

**Authors:** Joseph A.M. Smith, Lawrence J. Niles, Stephanie Feigin

## Abstract

Around the world, tidal flats play a unique ecological role in estuaries and are a primary feeding habitat for shorebirds and other benthic feeding organisms. Development and economic use of tidal flats can exclude species that depend on this habitat and disrupt ecological processes. In this study we examine patterns of abundance and feeding activity of American horseshoe crabs among oyster aquaculture structures on tidal flats that are adjacent to one of the most important horseshoe crab spawning sites in the world. We used custom-designed traps to sample horseshoe crab abundance beneath rack and bag aquaculture structures and adjacent areas without structures. In addition, we developed predictive spatial models representing three hypotheses regarding the movement of horseshoe crabs through aquaculture structures when transiting to and from spawning beaches. We tested the predictive power of each model using data from traps and found the strongest support for an avoidance model, where on average, horseshoe crabs are avoiding arrays of aquaculture structures when moving across inundated tidal flats. The best-supported spatial model also indicates that patterns of structure avoidance by horseshoe crabs can potentially affect abundance on spawning beaches, particularly with larger gear arrays that are closer to shore. We found additional support for aquaculture structure avoidance by examining an independent data set of horseshoe crab feeding pits on the tidal flats. Patterns of feeding pit density mirrored our trapping results, with fewer pits beneath and among aquaculture structures when compared to adjacent control areas. Horseshoe crabs are important constituents of the benthic food web and their displacement by aquaculture may translate to significant disruptions to the ecological function of tidal flats. This impact can be limited through deliberative spatial planning that seeks to balance ecological and economic management objectives.

## Introduction

Worldwide, tidal flats are an important ecological resource (Murray et al., 2019) that is being increasingly threatened by climate change (Passeri et al., 2015) and development (Song et al., 2013). While these forces drive the outright loss of tidal flats, other human uses such as harvest practices (Ferns et al., 2000) and aquaculture (Forrest et al., 2009) may affect the ecological composition and function of tidal flats.

Aquaculture activities may compete for space with organisms that depend on tidal flats such as shorebirds, benthic invertebrates and benthic-feeding animals. Research is needed to understand how species respond to aquaculture structures and tending activities to inform spatial planning of tidal flat use that is intended to meet both human and ecological needs. Without such an approach, development can potentially proceed to the detriment of ecological function.

In New Jersey a 600-acre tidal flat is situated at the epicenter of the world’s largest population of spawning horseshoe crabs and a globally important shorebird stopover. The expanse is ideal for intertidal aquaculture but important questions remain regarding the impact of aquaculture

Ideally the exclusion by intertidal oyster aquaculture of key ecological constituents is minimized and there is mutual use of the space. In this study we test this potentiality by examining patterns of horseshoe crab activity among oyster aquaculture structures and adjacent open-bottom areas. We use field data and spatial models which are validated with these data to examine a series of hypothesis corresponding with different potential effects of aquaculture on horseshoe crab movements. The results of this work are intended to help inform stakeholders, regulators and spatial planners in their efforts to balance economic and ecological objectives for the use and conservation of Delaware Bay tidal flats.

## Methods

### Study site

The study was conducted along a 1km shoreline segment occupied by intertidal structural oyster aquaculture of varying density and configurations. The intertidal flats have an undulating topography composed of lower sloughs and higher bars. Bars and some sloughs are exposed at low tide up to 400m offshore. At low tide horseshoe crabs tend to vacate bars and aggregate offshore or in water-filled sloughs. Rack structures are situated in both sloughs and bars, depending on the operation. Aquaculture rack arrays occupy approximately 1 Ha of area (including lanes between structures) and range from 105 to 340m offshore. Individual arrays range in size from (86-1,437 sq m). Aquaculture occurs in two high-density clusters separated by a gap of approximately 400m, although a few scattered smaller gear clusters occur in this intervening area.

Individual rack structures are made of rebar and vary in design but are typically 10 x 2.5’ with 8 supporting legs of varying height. Racks are placed end-to-end and are arrayed in double rows with approximately 1.5’of space between paired rows and 4-8’ spacing between double rows to make access lanes. Oysters in polyethylene bags rest on top of racks. Oysters are tended during low tide and are accessed with ATVs.

### Horseshoe crab sampling

To sample horseshoe crab activity on intertidal flats during periods of tidal inundation, we employed a custom trap design. Traps were constructed from high density polyethylene 1.5” mesh fencing. The fence was used to create a 6’ x 6’ x 4’ high open-topped holding pen around 6’ steel T-posts. On the down-bay side of the pen, a hinged one-way door (2’ wide x 1’ high) was constructed at the base of the fencing to allow horseshoe crabs in. A 7’ long, 1’ high lead fence was connected perpendicularly to the inshore edge of the door entrance and a smaller 1’ x 1’ baffle was connected perpendicularly to the opposite door. T-posts were also attached to the bottom of each side of the holding pen and the lead fence to minimize escapes under the fencing. All traps were constructed with the same design and orientation, with the concept that the door placement increased the probability of catching crabs on rising tides when currents are moving toward the door-side of traps. Horseshoe crabs are known to travel with prevailing tidal currents (Shuster and Botton, 1985).

Traps were systematically distributed among aquaculture structure arrays, with 8 total traps deployed. An additional 8 control traps were paired with traps in structures (6 traps in northern section of study area, 10 traps in the southern section). These were set nearby in a similar geomorphic and geographic setting (i.e. the same slough or bar). The lead fence was designed to be fitted beneath a double row of racks and span their width. This ensured that horseshoe crabs entering traps had to first pass beneath aquaculture structures. Rack heights above lead fences were 30cm for 6 traps and 23cm for 2 traps.

Traps set to catch crabs moving under structures were placed within arrays ranging from 228 and 1437.5 m^2^ (mean 790.4 m^2^) in size. Traps within arrays were set between 2.1 and 12.5 m from the array edge. Control traps were between 4.5 and 27m away (mean 14.4m) from adjacent aquaculture structures. 6 of 8 traps were set on the south side of arrays and 2 were set on the north side of arrays.

Once erected, traps were checked at least once daily during the diurnal low tide. All captured horseshoe crabs were counted and sexed. They were immediately released at a minimum distance of 25m in the next shoreward slough from the trap. Traps were checked over the course of 34 days between May 5 and June 8 2019 with a total of 560 individual trap checks.

### Trap data analysis

To estimate the overall difference between trap catches inside and outside of aquaculture structure arrays and patterns over space and time, we used repeated measures Generalized Estimating Equation models that were fit using the negative binomial distribution with a log link function and AR(1) working correlation structure. We chose this correlation structure because repeated measures show a decreasing correlation over time due to temporal variation horseshoe crab activtity. All analyses were conducted in SPSS software (IBM SPSS Statistics for Windows 2016). Estimated marginal means for categorical variables were contrasted with least significant difference post-hoc pairwise comparisons to test for differences among groups.

### Aerial mapping

To map the extent of aquaculture operations during the study, we collected aerial imagery of the study area (6/06/2019) using an DJI Mavic Pro drone programmed to collect imagery on a grid pattern at an altitude of 75m with 75% image front overlap and 65% image side overlap using DroneDeploy software (www.Dronedeploy.com). The mission collected 729 images which were compiled to create a seamless georeferenced orthomosaic image (2.54cm per-pixel resolution).

The resulting aerial image was used in GIS to develop spatial representations of aquaculture structures for subsequent use in spatial modelling. We also used this image for examining spatial patterns of horseshoe crab and ray feeding depressions on intertidal flats.

### Spatial modelling

We used Circuitscape software to develop spatial models representing three distinct hypotheses regarding horseshoe crab movement patterns among aquaculture structures. These were 1) no effect, 2) rack leg impedance and 3) structure avoidance.

Circuitscape uses circuit theory to model multiple movement pathways between nodes in a landscape (McRae et al., 2008) and landscape surfaces can be parameterized with varying resistance values to examine how movement may be affected by heterogeneous landscapes (McRae, 2006). Current maps generated by circuit theory algorithms are directly proportional to movements generated via random walks (Doyle and Snell, 2000).

We defined two focal regions (i.e. nodes) in order to simulate the movement of horseshoe crabs to and from spawning beaches. This included a linear offshore region that was a consistent distance from shore and an inshore region representing the footprint of spawning beaches along the shoreline. We then defined three resistance surfaces to represent the three hypotheses listed above.

The “no effect” hypothesis predicts no avoidance of structures by horseshoe crabs with movements unimpeded by rack structure legs. Crab abundance under this scenario is predicted to be equivalent inside and outside of aquaculture gear arrays. In Circuitscape, this is represented by a simple null model of movement between focal nodes with no variation in resistance across the study area.

The rack leg impedance hypothesis predicts no avoidance of structures by horseshoe crabs, but rack legs present an obstacle that requires crabs to redirect themselves once a rack leg is encountered. The frequent redirection necessitated by large numbers of obstacles would result in more time spent among aquaculture structures and, as a result, a higher observed abundance of crabs among structures compared to areas without structures. In Circuitscape, this is represented by locations with rack legs having infinite resistance with no other variation in resistance across the study area.

Finally, the avoidance scenario proposes that horseshoe crabs tend to avoid aquaculture structure arrays when moving to and from spawning beaches. In this case, the footprint of aquaculture structure arrays is represented initially in Circuitscape with double the resistance of the surrounding area. To develop an empirical basis for parametrizing this resistance value, we iteratively adjusted the resistance value representing aquaculture areas to maximize the correlation between trapping data and the resulting Circuitscape current map.

All focal and resistance surfaces were created at a 0.3 m resolution. Representations of aquaculture arrays were based on aerial imagery collected during the 2019 field season. To create the layer representing rack leg impedance, we initially created a point layer in GIS representing all rack legs in the study area. We then thinned the number of legs to half to ensure that the final resistance surface had evenly-spaced (1.5-2 m by 1m spacing) linear series of features representing rack arrays and adjacent lanes without clumping of features at the 0.3 m resolution.

We ran Circuitscape for each hypothesized scenario and generated a current map for each hypothesis using the Pairwise modeling mode and with movement from individual raster cells modelled based on a connection to eight neighboring cells. The resulting current density maps depict variation that is interpreted as the net movement probability or flow of random walkers through a region. This value can be used to predict passage rates and abundance of animals in the landscape.

We tested these predictions by relating current density maps to our horseshoe crab trap captures. For each hypothetical spatial model we extracted map values from the passage probability map at each trap location and examined the relationship between passage probability and horseshoe crab trapping data. The spatial model that best reflects reality would have the highest correlation with trapping data.

### Feeding depression mapping

Horseshoe crabs leave distinct feeding depressions in intertidal areas (Wan-Jean, 2010). Likewise, at our study site cownose rays also leave distinct feeding depressions (Howard et al., 1977; Smith and Merriner, 1985). Depressions are ephemeral, appearing whenever horseshoe crabs and rays are active in the study area, and disappearing over time or when high wind and wave action rework intertidal sediment. The two types of feeding pits vary by size, with the average pit diameter for horseshoe crabs being 20cm, while ray feeding depressions are considerably wider (up to 1m diameter) and deeper (Orth, 1975). We used aerial imagery to assess the distribution and abundance of feeding pits in our study area (Takeuchi and Tamaki, 2014). To compare the prevalence of feeding depressions beneath aquaculture structures and adjacent open areas, we delineated all structures in GIS that did not have oyster bags on top of them at the time our aerial imagery was taken. Such locations allow for a clear view under racks to determine the presence of feeding depressions. Most of these structures previously had bags on them during the study period. We then delineated adjacent control areas without aquaculture structures that were in the same geographic and geomorphic setting (i.e. similar distance from shore and occupying the same slough or bar). Control areas range from 0 to 25m from sampled rack areas. All delineated areas were fully exposed at low tide.

We generated 120 spatially-balanced random points (Theobald et al., 2007) in control and aquaculture structure areas and overlaid a regular 1m square grid across the study area. We then documented the presence and diameter of feeding depressions in all grid squares that contained a random point. We counted any depression that fell within, or intersected any boundary of the sample quad. We used normal mixture modelling to size-classify feeding depressions and used logistic regression to determine the probability of feeding depressions occurring within aquaculture structures and control areas. We also compared feeding depression occurrence between the northern and southern sections of the study area.

## Results

### Horseshoe crab sampling

Over the course of the study, 3,678 horseshoe crabs were captured comprising a male: female ratio of 2.3:1. This is similar to sex ratios observed in recent Delaware Bay trawl data (2:1, 95% CI 1.3-2.7) (Atlantic States Marine Fisheries Commission, 2019).

There was considerable variation in catch abundances over time, with the highest capture rates around the full moon in May followed by a decline in catch rates and a lower abundance spike during the subsequent new moon in June (fig 1a). There was a sudden drop in water temperature on 5/13 that appeared to cause trap catches to drop significantly for a day. Subsequently water temperatures increased and catches likewise increased.

**Figure 1.**
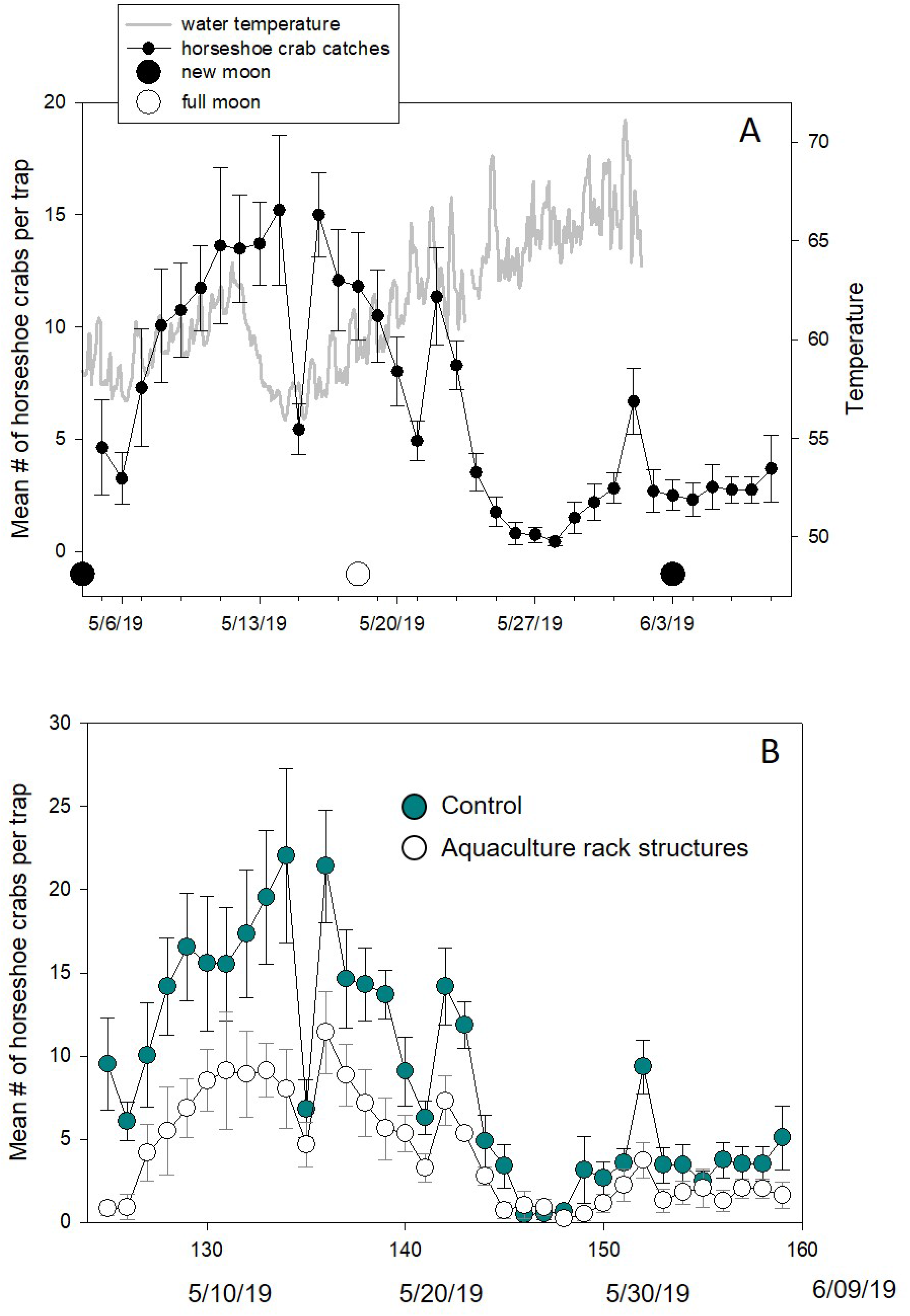
A) Overall patterns of horseshoe crab captures in traps reflecting predictable patterns of activity in relation to moon phase and water temperature; B) The same time sequence of trap captures separated by trap location within or outside of oyster aquaculture structure arrays.

We used a Generalized Estimating Equation model to examine variation in horseshoe crab trap catch abundance as function of date and a trap’s placement in control vs aquaculture structure areas, in northern vs southern section of the study area and its distance from shore.

Results of this model indicate a 48% lower catch abundance of horseshoe crabs beneath aquaculture structures (Wald χ2 = 22.03, d.f. = 1, p < 0.0001, Fig. 2a). The difference between control and aquaculture treatments was consistent over time across 34 daily trials (Fig 1b).

**Figure 2.**
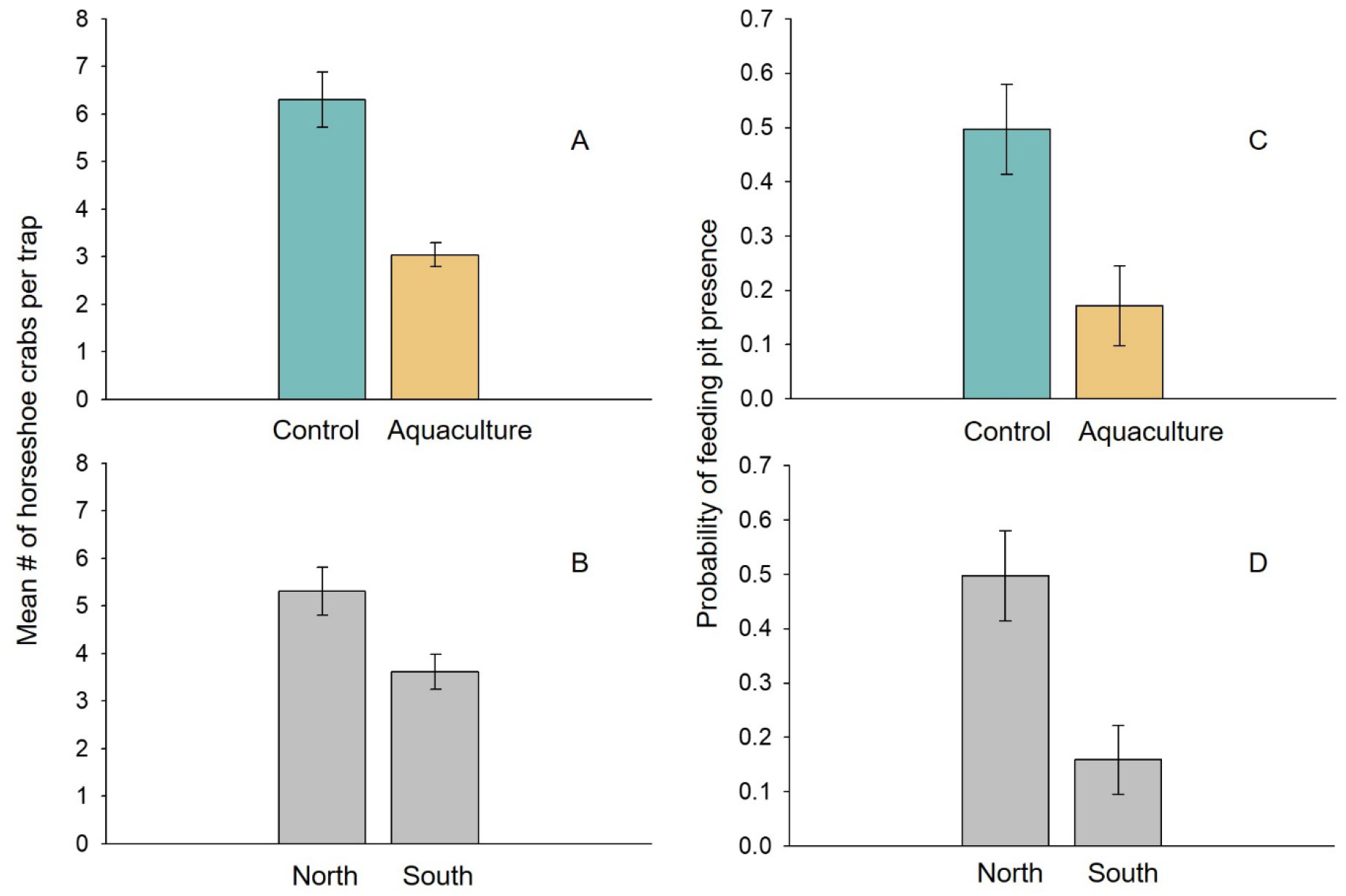
A) A comparison of mean capture rates of horseshoe crabs within or outside of oyster aquaculture structure arrays; B) A comparison of mean capture rates of horseshoe crabs in the northern vs southern regions of the study area; C) A comparison of mean encounter probability of horseshoe crab and ray feeding pits within or outside of oyster aquaculture structure arrays; D) A comparison of mean encounter probability of horseshoe crab and ray feeding pits in the northern vs southern regions of the study area.

The model also indicated higher catch abundances in the northern region of the study site (Wald χ2 = 14.59, d.f. = 1, p < 0.0001, Fig. 2b) while controlling for differences between catches beneath structures and control areas. Catches also varied with distance from shore, with relatively fewer horseshoe crabs caught offshore (Wald χ2 = 14.59, d.f. = 1, p =0.014).

To conduct a comparison of trap catch abundances for paired traps in aquaculture structures and adjacent control areas, a second Generalized Estimating Equation was constructed. This model included date, a “pair” variable denoting a trap’s pair membership, its position either within aquaculture structures or in a control area and an interaction term between pair membership and the structure/ control treatment. This model again indicated that traps set within aquaculture structures captured consistently fewer horseshoe crabs than in adjacent paired control traps in the same geographic and geomorphic setting (p<0.0001 for all 8 trap pair comparisons, Fig 3).

**Figure 3.**
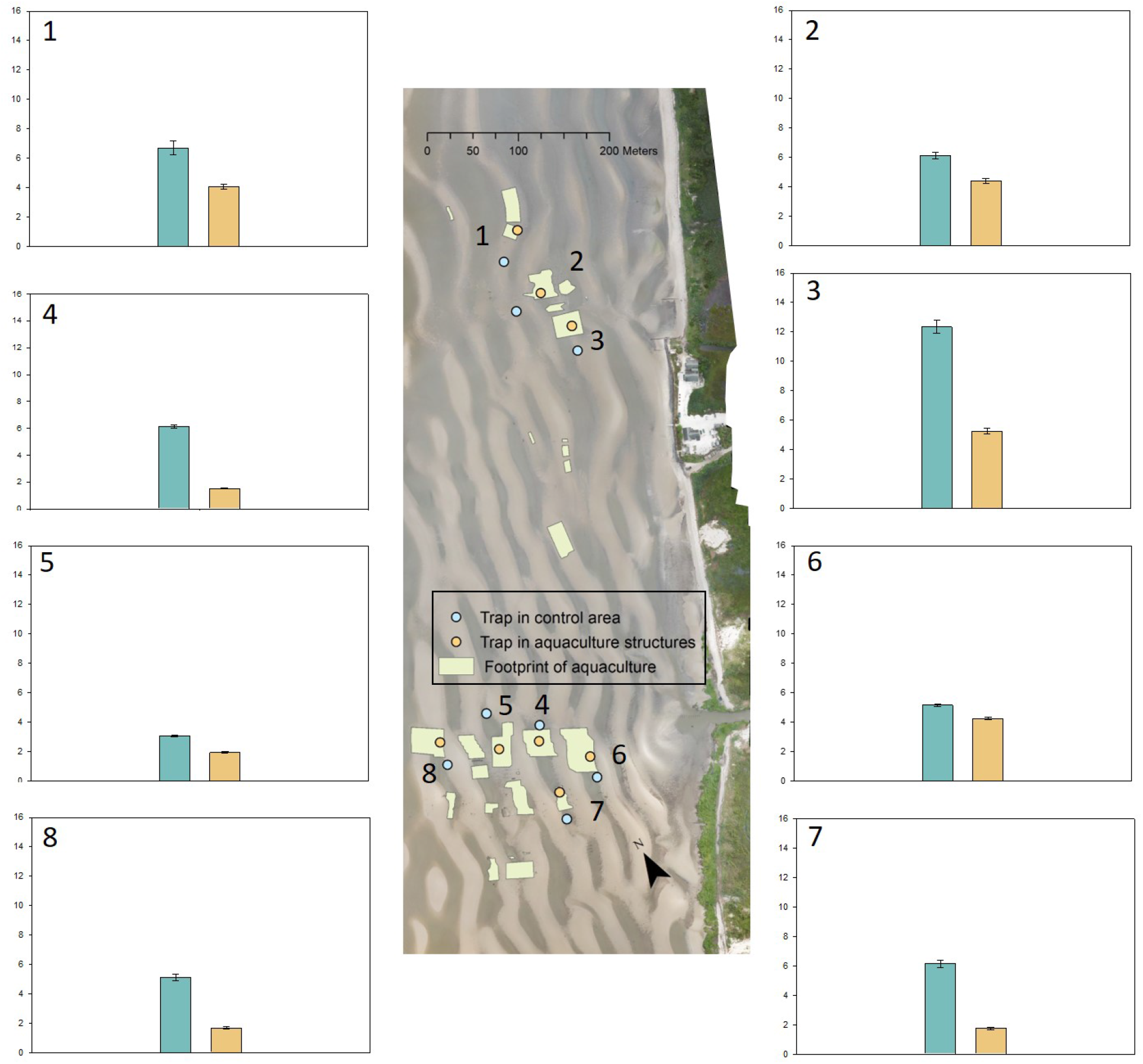
Pair-by-pair comparison of mean horseshoe crab capture rates of traps in the same locality within or outside of oyster aquaculture structure arrays. Y-axis represents mean capture rate. All difference between groups are significant.

### Spatial modelling

Maps depicting the relative probability of horseshoe crab passage generated for each hypothesis exhibited distinct spatial patterns. The null model, which hypothesized no effect of aquaculture structures on movement, displayed variation in current density from north to south as a result of variation in the coastline (Fig 4a). The shoreline in the northern part of the study area is is retreating at a slower rate than the southern part of the study area (0.9m/year in the north vs 2.5m/year in the south). The circuit-theory algorithm routes greater current density in areas where focal regions are closer together. This yielded greater current densities in regions exhibiting slower retreat.

**Figure 4.**
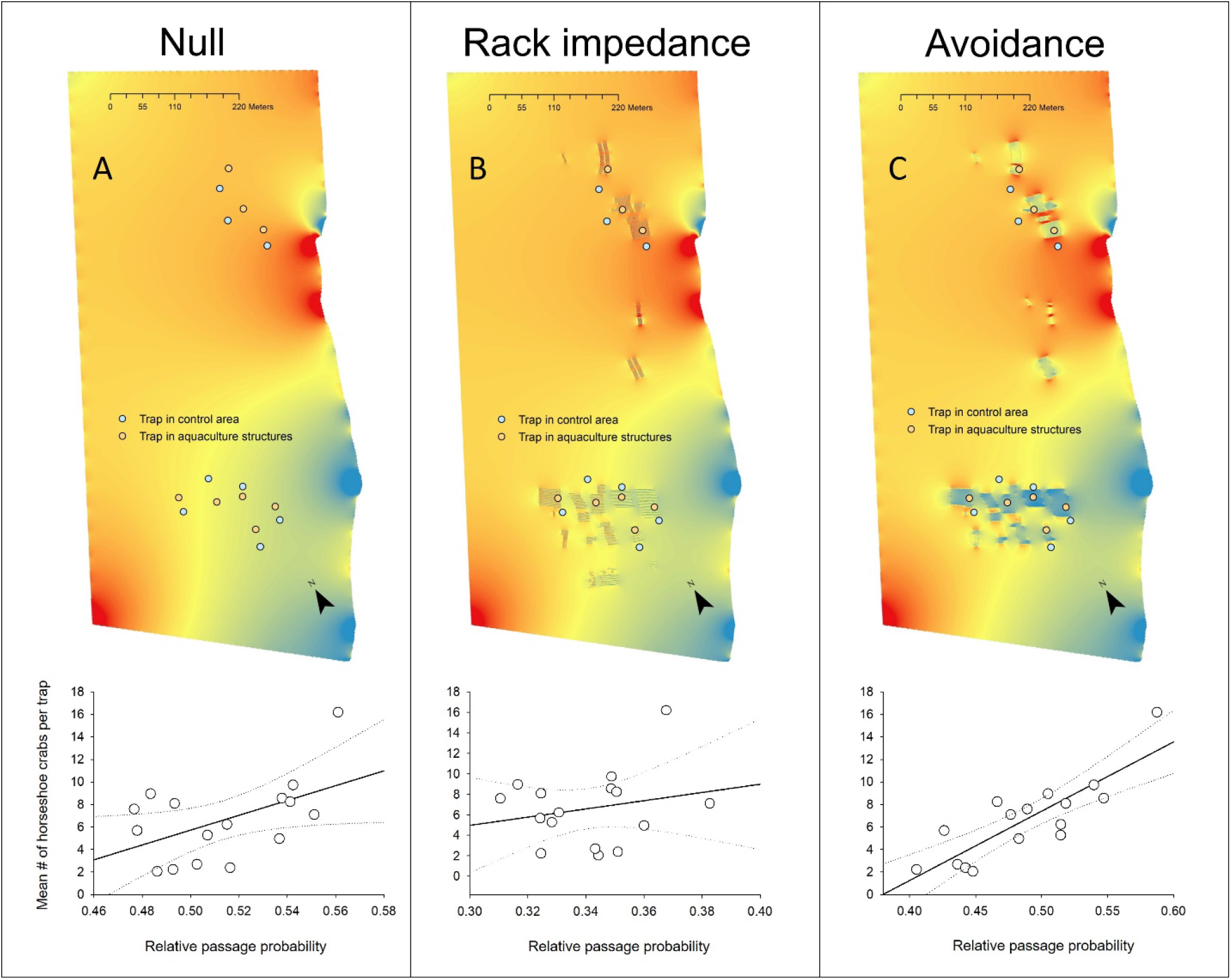
Upper panel: three spatial modelling representing different hypotheses regarding the effect of oyster aquaculture structures on horseshoe crab movements. The blue-to-red gradient represents varying levels of relative passage probability through a particular area, with blue being the lowest value and red the highest. Lower panel: map values are extracted for each trap location and regressed against the mean catch value for traps. The avoidance model shows the best correspondence with field data.

The rack leg impedance and avoidance models integrate the effect of the null model along with distinct effects of aquaculture structures. For the rack leg impedance hypothesis, the resulting current map indicated greater current densities within aquaculture structure arrays (Fig 4b). For the avoidance hypothesis, the resulting map indicated lower current densities within structure arrays, with higher densities in the region immediately surrounding arrays (Fig 4c). Both scenarios predict a “shadowing” effect on adjacent spawning beaches, where current densities are reduced compared with the null model (Figure 5).

**Figure 5.**
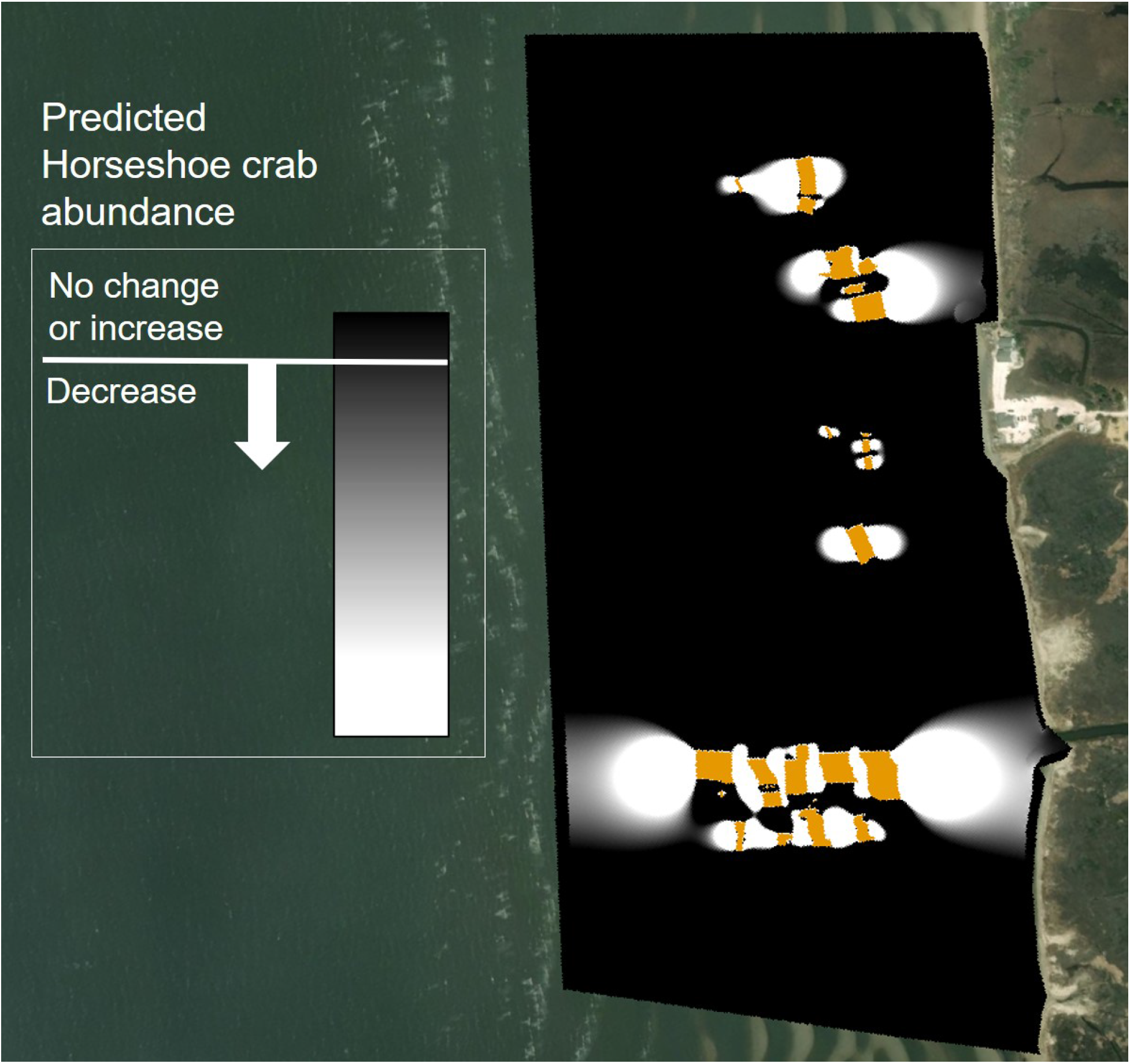
Predicted change in horseshoe crab densities as a result of aquaculture avoidance. The effect of rack avoidance is depicted, isolated from spatial patterns evident in the “no effect” model. This indicates that lower horseshoe crab densities are predicted inshore and offshore of gear arrays.

Regression analysis to test the predictive power of each current map indicated that the null model alone was a significant predictor of mean horseshoe crab catch abundances in traps (Fig 4a, F = 4.89, R^2^=0.26, df= 13, P=0.04, n=16) while there was no significant relationship between catch abundances and the rack impedance model (Fig 4b, F = 0.70, R^2^=0.05, df= 13, P=0.42, n=16). For the aquaculture structure avoidance model, the initial parameterization of arrays representing double the resistance of surrounding areas was significant and an improved fit compared to the null model (F = 21.07, R^2^=0.60, df= 13, P = 0.0004, n=16). Iterative adjustment of the resistance parameter to maximize the model R^2^ arrived at a resistance value that was 15% higher than the surrounding area (Fig 4c, F = 33.20, R^2^=0.70, df= 13, P <0.0001, n=16).

### Feeding depressions

A normal mixture model partitioned the feeding pit data into two size groups. We considered the smaller-diameter group to be putative horseshoe crab feeding pits (mean = 0.18m ± 0.02 SE) and the larger-diameter pits to be evidence of ray feeding (0.68m ± 0.05 SE). The smaller group mean values correspond well with published feeding depression estimates for horseshoe crabs (Wan-Jean, 2010) and the larger group corresponded well with that of cownose rays (Smith and Merriner, 1985).

Overall, 25 out of 120 sample quadrants contained at least one feeding depression. Five of these depressions were under aquaculture structures and 20 were in control areas outside of structures. None of the smaller, putative horseshoe crab feeding depressions were detected under aquaculture structures. A Generalized Linear Model including all feeding pit detections indicated significantly fewer feeding depressions overall under aquaculture structures when compared with control areas (Fig 1c, χ^2^ = 8.98, p = 0.0027). In addition, pits were more frequent in the northern portion of the study area when compared with the southern portion (Fig 1d, χ^2^ = 10.94, p = 0.0009).

## Discussion

These results show a clear pattern of reduced horseshoe crab activity beneath oyster aquaculture rack structures. Captures of horseshoe crabs in traps set among aquaculture structures caught significantly fewer crabs than control traps that were outside of structure arrays. This pattern was consistent over 34 daily trials across 8 paired replicates of control and aquaculture traps. An additional line of evidence derived from mapping horseshoe crab feeding pits also showed significantly reduced feeding activity of horseshoe crabs beneath oyster aquaculture structures.

A spatial model that depicted horseshoe crab avoidance of aquaculture structure arrays was strongly correlated with trapping results. In contrast, a model that predicted free movement of crabs under racks with the need for crabs to redirect after encountering rack legs showed no significant relationship with trapping data. A null model that predicted free movement of crabs through the study area with no effect of aquaculture structures did show a significant correlation with trapping data, but this relationship was weaker than the avoidance model.

The null model predicts a north-to-south trend in horseshoe crab passage probability that is driven by the shape of the coastline. Recessed, rapidly transgressing sections of the coast in the south show lower values while the northern section that is transgressing more slowly shows higher values. This north-to-south pattern correlates with the strong north-south difference we found between traps in the northern and southern clusters of aquaculture arrays. Likewise, horseshoe crab feeding pits were more prevalent in the northern cluster.

The difference in abundance from north to south could simply indicate that more shoreward regions of coastline promote aggregation of horseshoe crabs. Alternatively, shoreline position may serve as a proxy for habitat quality. The rapidly transgressing region of the shoreline to the south is also characterized by its poorer spawning habitat, with shallower beach sand depths and sections of exposed peat that are associated with reduce horseshoe crab spawning activity (Botton et al., 1988).

This pattern is validated by field sampling of horseshoe crab eggs on beaches in 2019 (unpublished data). This sampling records the depth of sand in sample pits to a threshold depth of 40cm that corresponds with high habitat quality (Smith et al., in press). In the northern beach segment of the study area, 100% of sample pits reached this depth. Moving south, 74% of sample pits reached this depth in the central beach segment while just 64% of sample pits reached this depth in the southern-most beach segment.

Taken together, our findings indicate that horseshoe crabs tend to areas occupied by oyster aquaculture structures. This conclusion is based on both the reduced abundance of horseshoe crabs under structures and the near-absence horseshoe crab feeding pits under structures. This suggests that those horseshoe crabs that move under structures may not engage in normal feeding activity once there.

The mechanism for rack structure avoidance is unclear, but our results suggest some possibilities. As illustrated in the rack impedance hypothesis, if horseshoe crabs were simply responding randomly to rack leg obstacles, they would tend to aggregate under aquaculture structures. This is because the travel time of an individual under structures would invariably slow down as the matrix of obstacles is navigated. Alternatively, as implied by the avoidance hypothesis spatial model (validated by field data), horseshoe crabs may be reacting non-randomly to obstacles and redirecting away from them. That is, when rack legs on the edge of arrays are encountered by horseshoe crabs, they may on average redirect away rather than into the array.

The intertidal flats at our study area have been examined by researchers for decades. Some of this work has documented the importance of these flats as a feeding resource for horseshoe crabs where they seek bivalves (Mark L. Botton, 1984) and the vast amount of sediment turnover that is driven by horseshoe crab activity (Kraeuter and Fegley, 1994). Field experiments showed the horseshoe crab feeding pressure dramatically affects the abundance and species composition of bivalves at this site (Mark L. Botton, 1984). Cownose and other benthic-feeding rays similarly are a major force in structuring benthic species composition and sediment reworking (Howard et al., 1977; Orth, 1975) as they too seek bivalves and other benthic prey to feed upon (Smith and Merriner, 1985).

The pits that remain after horseshoe crab and ray feeding experience increased particle deposition that enhances feeding opportunities for benthic-feeding organisms (Yager et al., 1993). This includes red knots and other shorebirds because feeding pits also concentrate horseshoe crab eggs on tidal flats.

Benthic bioturbation by horseshoe crabs, rays and other organisms is a critical and fundamental process that is an example of “ecosystem engineering” where a cascade of direct and indirect effects of this process serve to shape the ecosystem itself (Meysman et al., 2006). With this in mind, the exclusion of major drivers of bioturbation and apex predators in the benthic food web implies a significant disruption in the ecological function of tidal flats as a result of oyster aquaculture structures. This impact can be limited through deliberative spatial planning that seeks to balance ecological and economic management objectives.

## Acknowledgements

This work was funded by a grant to the Conserve Wildlife Foundation of New Jersey from the National Fish and Wildlife Foundation’s Atlantic Flyway Shorebird Initiative. We offer sincere thanks to the oyster growers who granted permission to conduct research on their aquaculture leases and cooperated with our work including Ned Gaine (who also contributed to study design and provided logistical advice and support), Lisa Calvo, Betsy Haskins, Brian Harman, Barney Hollinger, Shaughn Juckett and Joe Morrow. For assistance with fieldwork we thank John Callahan, Theo Diehl, Mike Kilpatrick, Mike Pellew and Reydson Rafael.

## Literature Cited

Atlantic States Marine Fisheries Commission, 2019. Horseshoe Crab Benchmark Stock Assessment. Atlantic States Marine Fisheries Commission.

Botton, Mark L., 1984. Diet and food preferences of the adult horseshoe crab Limulus polyphemus in Delaware Bay, New Jersey, USA. Mar. Biol. 81, 199–207. https://doi.org/10.1007/BF00393118

Botton, Mark L., 1984. The importance of predation by horseshoe crabs, Limulus polyphemus, to an intertidal sand flat community. Journal of Marine Research 42, 139–161. https://doi.org/10.1357/002224084788506086

Botton, M.L., Loveland, R.E., Jacobsen, T.R., 1988. Beach erosion and geochemical factors: influence on spawning success of horseshoe crabs (Limulus polyphemus) in Delaware Bay. Marine Biology 99, 325–332.

Doyle, P.G., Snell, J.L., 2000. Random walks and electric networks. arXiv preprint math/0001057.

Ferns, P.N., Rostron, D.M., Siman, H.Y., 2000. Effects of mechanical cockle harvesting on intertidal communities. Journal of Applied Ecology 37, 464–474. https://doi.org/10.1046/j.1365-2664.2000.00509.x

Forrest, B.M., Keeley, N.B., Hopkins, G.A., Webb, S.C., Clement, D.M., 2009. Bivalve aquaculture in estuaries: Review and synthesis of oyster cultivation effects. Aquaculture 298, 1–15. https://doi.org/10.1016/j.aquaculture.2009.09.032

Howard, J.D., Mayou, T.V., Heard, R.W., 1977. Biogenic sedimentary structures formed by rays. Journal of Sedimentary Research 47, 339–346. https://doi.org/10.1306/212F7167-2B24-11D7-8648000102C1865D

Kraeuter, J.N., Fegley, S.R., 1994. Vertical disturbance of sediments by horseshoe crabs (Limulus polyphemus) during their spawning season. Estuaries 17, 288–294. https://doi.org/10.2307/1352578

McRae, B.H., 2006. Isolation by resistance. Evolution 60, 1551–1561.

McRae, B.H., Dickson, B.G., Keitt, T.H., Shah, V.B., 2008. Using Circuit Theory to Model Connectivity in Ecology, Evolution, and Conservation. Ecology 89, 2712–2724. https://doi.org/10.1890/07-1861.1

Meysman, F.J., Middelburg, J.J., Heip, C.H., 2006. Bioturbation: a fresh look at Darwin’s last idea. Trends in Ecology & Evolution 21, 688–695.

Murray, N.J., Phinn, S.R., DeWitt, M., Ferrari, R., Johnston, R., Lyons, M.B., Clinton, N., Thau, D., Fuller, R.A., 2019. The global distribution and trajectory of tidal flats. Nature 565, 222–225. https://doi.org/10.1038/s41586-018-0805-8

Orth, R.J., 1975. Destruction of eelgrass, Zostera marina, by the cownose ray, Rhinoptera bonasus, in the Chesapeake Bay. Chesapeake Science 16, 205–208. https://doi.org/10.2307/1350896

Passeri, D.L., Hagen, S.C., Medeiros, S.C., Bilskie, M.V., Alizad, K., Wang, D., 2015. The dynamic effects of sea level rise on low-gradient coastal landscapes: A review. Earth’s Future 3, 159–181.

Shuster, C.N., Botton, M.L., 1985. A contribution to the population biology of horseshoe crabs,Limulus polyphemus (L.), in Delaware Bay. Estuaries 8, 363–372.

Smith, J.A.M., Niles, L., Modjeski, A., Hafner, S., Dillingham, T., in press. Beach restoration improves habitat quality for American horseshoe crabs and shorebirds in the Delaware Bay, USA. Marine Ecology Progress Series.

Smith, J.W., Merriner, J.V., 1985. Food habits and feeding behavior of the cownose ray,Rhinoptera bonasus, in lower Chesapeake Bay. Estuaries 8, 305–310. https://doi.org/10.2307/1351491

Song, D., Wang, X.H., Zhu, X., Bao, X., 2013. Modeling studies of the far-field effects of tidal flat reclamation on tidal dynamics in the East China Seas. Estuarine, Coastal and Shelf Science 133, 147–160. https://doi.org/10.1016/j.ecss.2013.08.023

Takeuchi, S., Tamaki, A., 2014. Assessment of benthic disturbance associated with stingray foraging for ghost shrimp by aerial survey over an intertidal sandflat. Continental Shelf Research 84, 139–157. https://doi.org/10.1016/j.csr.2014.05.007

Theobald, D.M., Stevens, D.L., White, D., Urquhart, N.S., Olsen, A.R., Norman, J.B., 2007. Using GIS to generate spatially balanced random survey designs for natural resource applications. Environmental Management 40, 134–146.

Wan-Jean, L.E.E., 2010. Intensive use of an intertidal mudflat by foraging adult American horseshoe crabs Limulus polyphemus in the Great Bay estuary, New Hampshire. Current Zoology 5, 011.

Yager, P.L., Nowell, A.R.M., Jumars, P.A., 1993. Enhanced deposition to pits: A local food source for benthos [WWW Document]. https://doi.org/info:doi/10.1357/0022240933223819

